# ExplaiNN: interpretable and transparent neural networks for genomics

**DOI:** 10.1101/2022.05.20.492818

**Authors:** Gherman Novakovsky, Oriol Fornes, Manu Saraswat, Sara Mostafavi, Wyeth W. Wasserman

**Affiliations:** Centre for Molecular Medicine and Therapeutics, Department of Medical Genetics, BC Children’s Hospital Research Institute, University of British Columbia, Vancouver, BC V5Z 4H4, Canada; Division of Computational Genomics and Systems Genetics, German Cancer Research Center (DKFZ), Heidelberg, Germany; European Molecular Biology Laboratory (EMBL), Genome Biology Unit, Heidelberg, Germany; Paul G. Allen School of Computer Science and Engineering, University of Washington (UW), Seattle, USA

## Abstract

Sequence-based deep learning models, particularly convolutional neural networks (CNNs), have shown superior performance on a wide range of genomic tasks. A key limitation of these models is the lack of interpretability, slowing down their adoption by the genomics community. Current approaches to model interpretation do not readily reveal how a model makes predictions, can be computationally intensive, and depend on the implemented architecture. Here, we introduce ExplaiNN, an adaptation of neural additive models[1] for genomic tasks wherein predictions are computed as a linear combination of multiple independent CNNs, each consisting of a single convolutional filter and fully connected layers. This approach brings together the expressiveness of CNNs with the interpretability of linear models, providing global (cell state level) as well as local (individual sequence level) biological insights into the data. We use ExplaiNN to predict transcription factor (TF) binding and chromatin accessibility states, demonstrating performance levels comparable to state-of-the-art methods, while providing a transparent view of the model’s predictions in a straightforward manner. Applied to *de novo* motif discovery, ExplaiNN identifies equivalent motifs to those obtained from specialized algorithms across a range of datasets. Finally, we present ExplaiNN as a plug-and-play platform in which pretrained TF binding models and annotated position weight matrices from reference databases can be easily combined. We expect that ExplaiNN will accelerate the adoption of deep learning by biological domain experts in their daily genomic sequence analyses.

## Introduction

High-throughput genomics methods such as ATAC-seq[2] or ChIP-seq[3], which respectively assess genome-wide accessibility of chromatin and binding of transcription factors (TFs), allow functional annotation of DNA elements. The sheer scale of the data generated by these methods precludes manual analyses. Machine and deep learning have become pervasive in large-scale genomic analyses due to their ability to identify meaningful features in massive datasets (reviewed in [4,5]). For instance, deep learning models have shown superior performance in predicting genome folding[6], chromatin accessibility states[7–10], gene expression levels[11–13], and TF binding sites (TFBSs; reviewed in [14]).

While increasingly complex models are now feasible, some are opaque; they do not readily reveal the features and properties that underlie their predictions[15]. For genomics, there are multiple approaches to improve the interpretability of deep learning models (reviewed in [16]), of which we focus on filter visualization and attribution methods.

For convolutional neural networks (CNNs) and related models, a powerful interpretation approach is to visualize the filters of the first convolutional layer as position weight matrices (PWMs), a common format in bioinformatics to represent TFBS patterns[17], by separately aligning the set of sequences activating each filter[18]. The resulting filter PWMs can be assigned a biological annotation by comparing them to TF binding profiles from reference databases like JASPAR[19]. While filter visualization provides an overview of the genomic features that the model has learned in the first layer, the success of this approach is highly dependent on model architectural choices such as max pooling or filter sizes[20]. Moreover, it is difficult to interpret how the model combines the learned motifs within the deeper layers. Furthermore, to gain interpretability at a global level (*i*.*e*. how each filter influences the model’s predictions), the filters must be nullified sequentially, which is both computationally intensive and dependent on arbitrary thresholds[9,10].

Attribution methods quantify the importances of individual nucleotides in the input sequences using forward-(*e*.*g. in silico* mutagenesis[7,21]) or back-propagation[22,23]. These importance scores can be the basis for further clustering of activating sub-sequences into PWMs[24], which, as with filter visualization, can in turn be annotated biologically by comparing them to known TF motifs. Although attribution methods provide local interpretability by identifying important nucleotides in the input sequences, quantifying the contribution of each feature to the overall model’s predictions (*i*.*e*. global interpretability) remains challenging[25]. Noteworthy, neither approach is transparent as to how the model makes predictions.

Unlike deep learning models, linear models are interpretable and transparent: the basis on which they make predictions can be readily understood by evaluating the learned feature coefficients. Agarwal and colleagues recently introduced neural additive models (NAMs), which combine features of deep learning and linear models[1]. NAMs compute predictions as a linear combination of outputs from independent deep learning models, each tuned to one input feature, resulting in the levels of explainability much appreciated in linear models without compromising on accuracy.

In this study, we present ExplaiNN (explainable neural networks), a fully interpretable and transparent, sequence-based deep learning model for genomic tasks inspired by NAMs. We evaluate ExplaiNN on different tasks, demonstrating that it performs as well as state-of-the-art models while yielding similar interpretations to more complex approaches, both locally and globally, but in less time and in a more intuitive and simple manner. Next, we show that the motifs learned by the convolutional filters of ExplaiNN are equivalent to those discovered by *de novo* tools on the same data. Finally, we apply ExplaiNN as a plug-and-play platform for JASPAR PWMs and pre-trained deep learning models with which to interpret genomic data, such as by distinguishing the individual roles of TFs with highly similar DNA-binding specificities.

## Results

### ExplaiNN is a glass box deep learning model for genomics

ExplaiNN is a deep learning approach for genomic tasks trained on one-hot encoded sequences inspired by NAMs[1]. Predictions are computed as a linear combination of multiple independent CNNs (hereafter referred to as units), each consisting of one convolutional layer with a single filter followed by exponential activation, which has been seen to improve the motif representations learnt by CNN filters[26], and two fully connected layers (**Fig. 1A**). ExplaiNN provides global interpretability, as the filter of each unit can be mapped to a TF profile from JASPAR using Tomtom[27] (**Methods**), thus assigning a biological annotation to that unit. The main difference between ExplaiNN and a standard CNN is that the learned weights of each unit from the final linear layer (*i*.*e*. the coefficients) are interpretable, similarly to a linear model. In addition, ExplaiNN can also provide local interpretability by multiplying the output of each unit by the weight of that unit for each input sequence (hereafter referred to as unit importance scores; **Methods**).

**Fig. 1:**
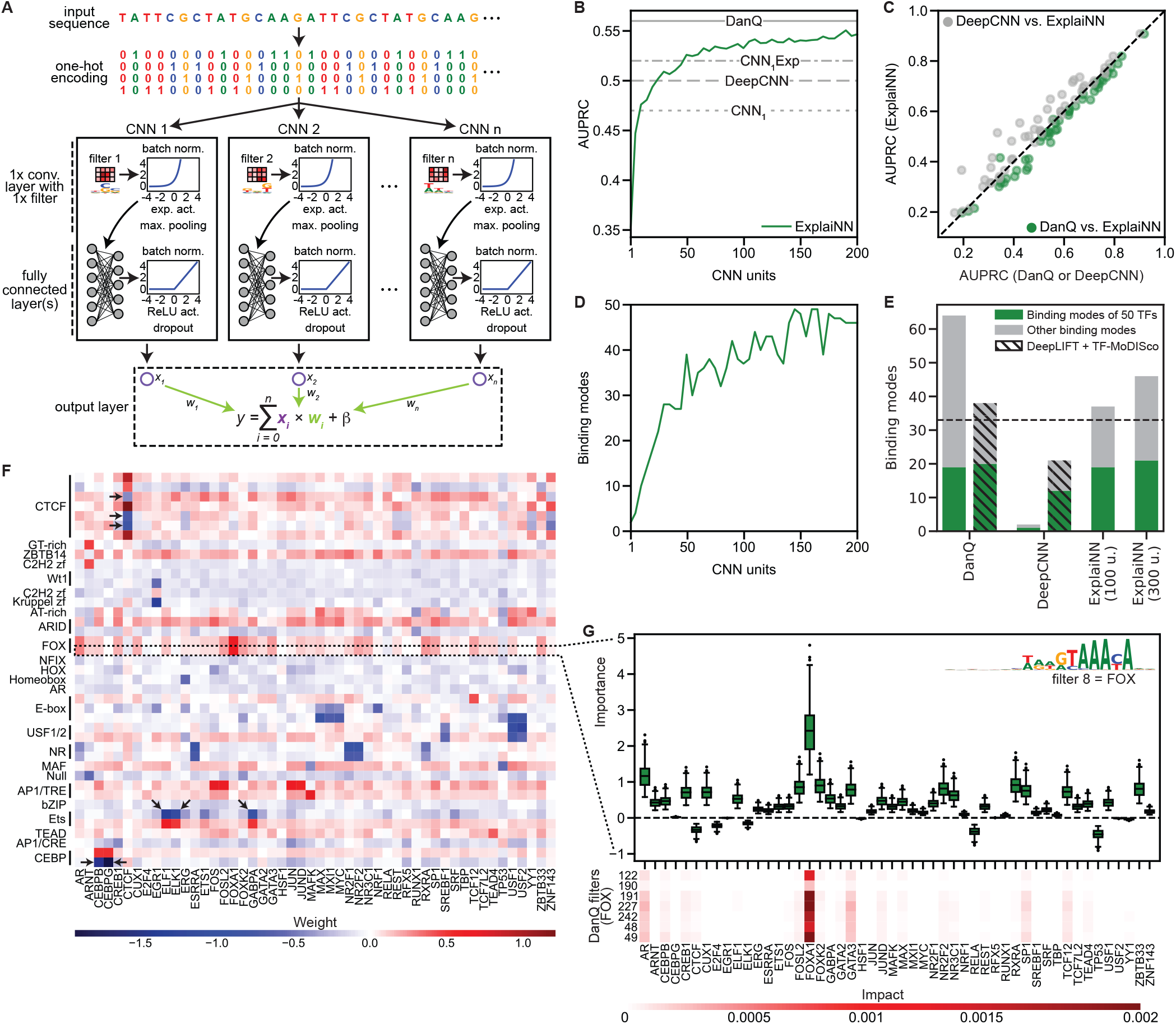
The ExplaiNN model and its application to TFBS prediction in OCRs. (**A**) ExplaiNN takes as input one-hot encoded DNA sequences. Architecturally, it is composed of multiple independent CNNs (*i*.*e*. units), each of which comprising one convolutional layer with a single filter, batch normalization, exponential activation, and max pooling, and two fully connected layers with batch normalization, ReLU activation, and dropout. The final linear layer of ExplaiNN (*i*.*e*. the output) combines the outputs from each unit (denoted here using Xs). (**B**) Performances (AUPRC; y-axis) of ExplaiNN models trained using increasing numbers of units (x-axis) on predicting the binding of 50 TFs in OCRs (green line). The performances of DanQ[8], a deep CNN with 3 convolutional layers (*i*.*e*. DeepCNN), and two shallow CNNs with 1 convolutional layer (*i*.*e*. CNN_1_ and CNN_1_Exp featuring an exponential activation function instead of ReLU) are provided as baselines (gray lines). (**C**) Pairwise comparison of the individual performances (AUPRC) of the 50 TFs from the previous dataset between the ExplaiNN model trained using 100 units and either DanQ (green dots) or the DeepCNN (gray dots). (**D**) Number of binding modes (y-axis) detected with ExplaiNN using increasing numbers of units (x-axis). (**E**) Number of binding modes detected with DanQ, the DeepCNN, and ExplaiNN trained using 100 and 300 units on the previous dataset using either filter visualization (no pinstripes) or TF-MoDISco[24] clustering on DeepLIFT[23] attribution scores (pinstripes). The 50 TFs in the dataset are represented by 33 unique binding modes (dashed line), some of which are detected using different combinations of models and interpretive approaches (green); other detected binding modes (*i*.*e*. different from those 33) are shown in grey. (**F**) Heatmap of the final linear layer weights of the ExplaiNN model trained using 100 units, with rows representing units with assigned biological annotations based on their Tomtom[27] similarity to known TF profiles from the JASPAR database[19] and columns representing the 50 TFs predicted by the model. More than one filter can learn the same TF motif representation, but some may not contribute to the model’s predictions (black arrows). (**G**) (top) Visualization of importance scores for a unit annotated as FOXA1 from the ExplaiNN model trained using 100 units. This unit contributes the most to the prediction of FOXA1 binding. (bottom) Filter nullification analysis for the seven DanQ filters annotated as FOX TFs. The results are consistent with the unit importance scores. AUPRC, area under the precision-recall curve; CNN, convolutional neural network; OCR, open chromatin region; ReLU, rectified linear unit; TF, transcription factor; TFBS, TF binding site.

As a proof of concept, we trained ExplaiNN to predict the binding of 50 TFs to open chromatin regions (OCRs) from a reference dataset describing the binding of 163 TFs to >1.8M 200-bp long OCRs that we repurposed for this study[28] (**Methods**). A key hyperparameter in ExplaiNN is the number of independent units to be used. To assess the impact of this hyperparameter on model performance, we trained multiple ExplaiNN models using increasing numbers of units (from 1 to 200). As expected, the performance of ExplaiNN improved with the number of units used, plateauing at around 100 units (**Fig. 1B**). For comparison, we evaluated four additional models on the same dataset (**Methods**): a CNN with one convolutional layer (CNN_1_); a CNN_1_ with exponential activation function (CNN_1_Exp); a deep CNN with three convolutional layers (DeepCNN); and DanQ[8], a hybrid deep learning model with convolutional and recurrent layers. ExplaiNN outperformed all three CNN models as measured by average area under the precision-recall curve (AUPRC) and, when using more than 100 units, nearly reached the performance of the more complex DanQ (**Fig. 1B**). Focusing on the ExplaiNN model trained with 100 units, it outperformed the DeepCNN model for most TFs, performing only slightly worse than DanQ (**Fig. 1C)**.

Next, we visualized the filters of each ExplaiNN model and assigned them TF binding modes from JASPAR, which we defined based on the hierarchically clustered groups of DNA-binding profiles included in the database (**Table S1**; **Methods**). As with performance, the number of binding modes recovered by the model increased with the number of units used (**Fig. 1D**). For comparison, we provide the number of binding modes recovered by the DeepCNN and DanQ when applying filter visualization, as we did with ExplaiNN, or when using TF-MoDISco[24] clustering on DeepLIFT[23] attribution scores (**Methods**). Out of 33 different binding modes associated with the set of 50 TFs analyzed, ExplaiNN models trained with 100 and 300 units recovered 19 and 21, respectively, which is a similar number as DanQ (19 when applying filter visualization and 20 when using DeepLIFT and TF-MoDISco) and greater than obtained for the DeepCNN model (**Fig. 1E**).

An advantageous feature of ExplaiNN is that one can readily visualize the final linear layer weights for global interpretation purposes (**Fig. 1F**; **Methods**). For example, units with filters annotated as FOX motifs had high positive weights for predicting the FOXA1 class. Similarly, CEBP-, CTCF-, and Ets-like units had high positive weights associated with predicting the classes of CEBP factors, CTCF, and Ets family members, respectively. However, some units had negative weights for predicting the class of their annotated TF (**Fig. 1F**; highlighted with arrows). To delve further into the contribution of each unit to the prediction of each class, we computed unit importance scores (**Methods**). Visualization of the importance scores of a FOX-like unit in a heatmap confirmed its importance for predicting the FOXA1 and AR classes (**Fig. 1G**; top panel), in agreement with the observation that FOXA1 helps shape AR signaling in prostate cells[29]. Visualizing unit importance scores also revealed an aspect as to why several CTCF-like units had negative weights for predicting the CTCF class: the importances of these units for the CTCF class were negligible, suggesting that the model was not using them to make predictions for that class (**Fig. S1**). The same was true for units annotated as CEBP and Ets with negative weights for those classes (**Fig. S1**). Finally, we calculated the impact from nullifying the FOX-like filters in the DanQ model one at a time (**Fig. 1G**; bottom panel; **Methods**) and, as expected, the impact scores from the filter nullification process were consistent with the unit importance scores.

Taken together, these analyses demonstrated that ExplaiNN performs comparably to more complex models, at least for TF binding prediction. In addition, ExplaiNN provided local and global interpretation quickly and readily compared to using DeepLIFT followed by TF-MoDISco or filter visualization and nullification.

### ExplaiNN captures homotypic but not heterotypic cooperative interactions

Given the architecture of ExplaiNN, in which each unit filter is independent of the rest, we expected that it would not be able to learn nonlinear interactions such as heterotypic cooperativity between pairs of TF motifs. In contrast, DeepSTARR is a CNN trained on STARR-seq data to predict the activities of *Drosophila* developmental and housekeeping enhancers that can capture these types of interactions[30]. For each pair of motifs analyzed, the authors of DeepSTARR performed a distance dependence analysis by sliding one motif along randomly generated sequences in the centre of which they embedded the second motif. Accounting for additive effects, they observed that the output of DeepSTARR increased when the two motifs were proximal, suggesting that the model had learned heterotypic cooperative interactions. To check whether this was also the case for ExplaiNN, we trained it on the same dataset and compared its performance to DeepSTARR by calculating the Pearson correlation coefficient (PCC) between predicted and actual enhancer activities (**Methods**). ExplaiNN performed worse than DeepSTARR on both developmental (PCC = 0.61 vs. 0.68) and housekeeping (PCC = 0.71 vs. 0.74) enhancers, which we attributed to the greater presence of nonlinear interactions in this particular dataset. Next, following the specifications from DeepSTARR, we performed a distance dependence analysis between the housekeeping TFs Dref, Ohler1, and Ohler6 (**Methods**). As a negative control, we slid the 5-mer GGGCT. As expected, DeepSTARR was able to learn distance dependencies between the three motifs (**Fig. 2**). This was not the case for ExplaiNN: during the analysis, the resulting model outputs using the three motifs were similar to sliding GGGCT (**Fig. 2**). While these results were expected given the architecture of ExplaiNN, we wondered if the fully connected layers would be able to capture homotypic cooperative interactions between pairs of instances of the same TF motif. We repeated the previous distance dependent analysis but this time sliding a second instance of the motif that had been fixed at the centre of the sequences. We observed that both ExplaiNN and DeepSTARR consistently captured homotypic cooperativity for Dref and Ohler6, however, for Ohler1, the distance dependencies obtained from the two methods were non-concordant (**Fig. 2**). To confirm whether the fully connected layers were responsible for learning these interactions, we trained a second ExplaiNN model on the DeepSTARR dataset in which the fully connected layers of each unit had been replaced with a global max pooling layer, after which we repeated the distance dependencies analysis. This model without fully connected layers was unable to learn any distance dependencies: results were similar to the negative control (**Fig. S2**). Taken together, and as we had anticipated based on its architecture, ExplaiNN is not suitable for detecting heterotypic cooperative interactions, however, the fully connected layers within each unit allow homotypic cooperativity to be learned.

**Fig. 2:**
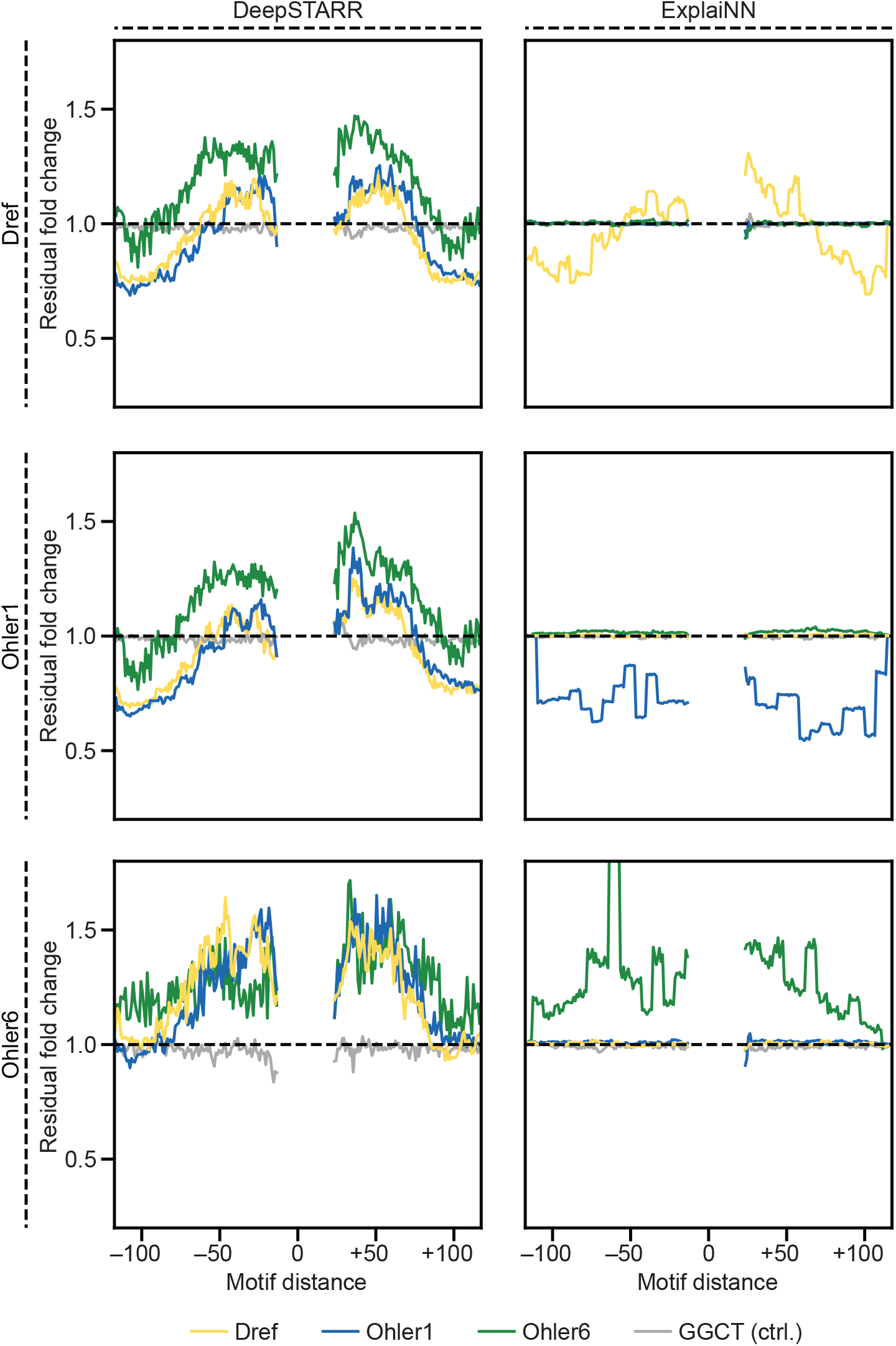
Exploring the limitations of ExplaiNN in capturing nonlinear interactions. Cooperativity (residual fold change; y-axis) plotted as a function of distance (x-axis) between the motifs of the housekeeping TFs Dref (top row; yellow), Ohler1 (middle row; blue), and Ohler6 (bottom row; green) for DeepSTARR[30] (left column) and ExplaiNN (right column). The 5-mer GGGCT is provided as a negative control (light gray). TF, transcription factor.

### ExplaiNN learns high-quality motifs comparable to *de novo* motif discovery tools

*De novo* motif discovery methods continue to emerge and improve[31–36]. With the dramatic escalation in the size of datasets, the execution time of these methods is increasingly a limitation. Furthermore, many *de novo* motif discovery methods are assay-specific, as exemplified by the DREAM5 challenge evaluation on protein binding microarray (PBM) data[37], requiring an extensive adjustment of parameters for their application to different assays. We sought to explore the capacity of ExplaiNN for efficient *de novo* motif discovery within a unified platform. For each of the 163 TFs from the previous dataset, we trained a model with 100 units and then visualized the filters and importance scores of each unit, resulting in 100 PWMs for each TF (**Methods**). As expected, PWMs derived from visualizing filters associated with important units performed better: for 139 TFs (85.3%), the best performing PWM was derived from the filter of one of the 10 most important units (**Fig. 3A**). Next, we applied STREME[34], a state-of-the-art method for *de novo* motif discovery, on the same sequences used for training the ExplaiNN models to discover 100 motifs for each TF (**Methods**). Pairwise comparison of the performances of the best PWMs obtained by each method revealed subtle differences (**Fig. 3B**), although PWMs discovered by STREME were superior for TFs with small training datasets (**Fig. 3C**). Notably, the execution times exhibited by the two methods differed greatly, with ExplaiNN being >100 times faster for TFs with large training datasets of ≥50,000 sequences (**Fig. 3D**). These differences were consistent with recent reports related to the benefits of GPU-enabled *de novo* motif discovery[33,36].

**Fig. 3:**
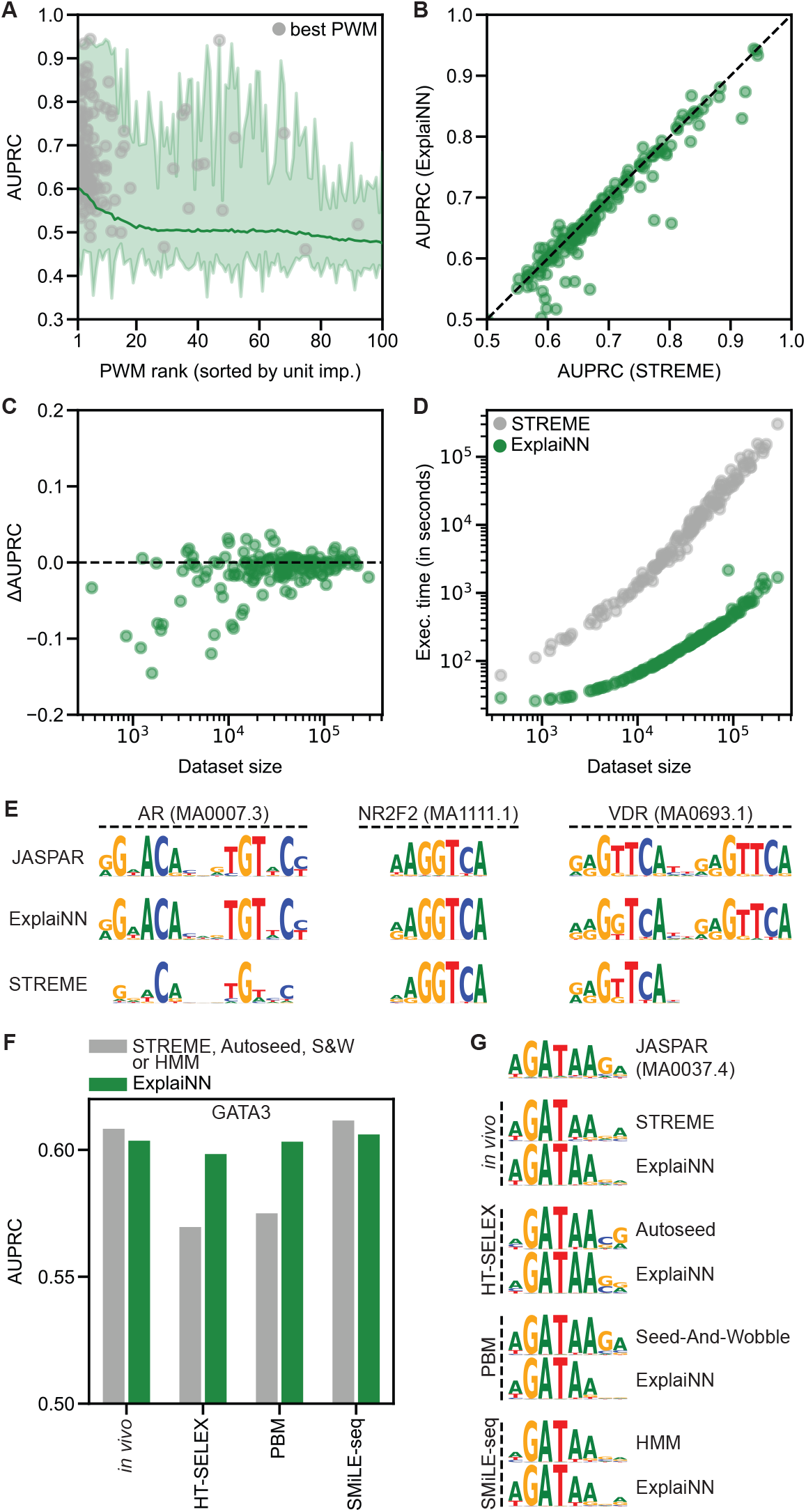
*De novo* motif discovery with ExplaiNN. (**A**) Average performances (AUPRC; y-axis) by rank (x-axis) of PWMs derived from training ExplaiNN models with 100 units on *in vivo* datasets of 163 TFs and then visualizing the filter of each unit (*i*.*e*. 100 PWMs per TF). The rank of each PWM is given by the importance of its unit. The gray dots indicate the rank of the best performing PWM for each TF. (**B**) Pairwise comparison of the individual performances (AUPRC) of the best PWMs derived for each TF from the previous dataset using ExplaiNN (y-axis) or STREME[34] (x-axis). (**C**) Performance difference (*i*.*e*. ΔAUPRC) of the previous PWMs (x-axis) is plotted with respect to the dataset size of the corresponding TF (x-axis). (**D**) Execution time (in seconds; y-axis) of the *de novo* motif discovery application of ExplaiNN (green dots) and STREME (gray dots) is plotted with respect to the dataset size of the corresponding TF (x-axis). (**E**) Logos derived using ExplaiNN or STREME for the nuclear receptors AR, NR2F2, and VDR from the previous dataset. For comparison, the JASPAR[19] logos for these TF profiles are shown: MA0007.3 (AR), MA1111.1 (NR2F2), and MA0693.1 (VDR). (**F**) Performances (AUPRC; y-axis) of PWMs derived from different experimental assay datasets related to the TF GATA3 by different methods (x-axis), including ExplaiNN (green bars) and four assay-specific methods[34,41–43] (grey bars). (**G**) GATA3 logos derived from the dataset of each experimental assay using ExplaiNN or the assay-specific method. The JASPAR logo for this TF profile (MA0037.4), derived by applying RSAT[35] on the mouse Gata3 ChIP-seq data from ReMap[44], is shown at the top. AUPRC, area under the precision-recall curve; HMM, hidden Markov model; PBM, protein binding microarray; PWM, position weight matrix; S&W, Seed-And-Wobble; TF, transcription factor.

The convolutional filters of CNNs are not guaranteed to learn full motif representations. Instead, CNNs are designed to learn motifs in a hierarchical manner, with the filters learning partial motifs that are assembled into full motifs in deeper layers, such as through higher-order interactions facilitated by the fully connected layers[20]. To explore the ability of ExplaiNN to learn full motif representations, we considered the case of nuclear receptors, which can bind to DNA sequences in the form of palindromes, direct repeats, or extended monomeric sites[38]. Focusing on 12 nuclear receptors from the previous analysis with at least 10,000 training sequences, ExplaiNN was able to learn motif representations faithful to the profiles of those TFs from the JASPAR database, while STREME often reported partial motifs consistent with monomeric sites (**Fig. 3E** and **S3**).

Motivated by the success of ExplaiNN in discovering motifs in *in vivo* data, and to demonstrate its capacity on data from various assays, we benchmarked ExplaiNN against other *de novo* methods, but this time using *in vitro* data (**Methods**). We downloaded publicly available HT-SELEX[39], PBM[40], and SMiLE-seq[41] data for the TF GATA3, as well as the PWMs discovered in these datasets by three assay-specific *de novo* motif discovery methods: Autoseed[42] (HT-SELEX), Seed-and-Wobble[43] (PBM), and a hidden Markov model (HMM) approach (SMiLE-seq). The performance of the best PWMs derived with ExplaiNN was consistent regardless of the type of assay (**Fig. 3F**), as were their logos (**Fig. 3G**). Focusing on specific assays, the capacity of ExplaiNN to discover *de novo* motifs in the *in vivo* and SMiLE-seq data, as measured by the performance of the best PWMs derived, was comparable to that of STREME and the HMM-based method, respectively, while outperforming Autoseed and Seed-and-Wobble in their corresponding assays (**Fig. 3F**). All GATA3 logos were very similar to each other, and were also similar to the logo from JASPAR for this TF profile (MA0037.4), derived originally by applying RSAT[35] on mouse Gata3 ChIP-seq data from ReMap[44] (**Fig. 3G**). Taken together, this supports the potential of ExplaiNN as a fast, universal method for *de novo* motif discovery.

### ExplaiNN recapitulates the *cis*-regulatory lexicon of immune cell differentiation

To further explore the capabilities of ExplaiNN on distinct data types, we compared its performance against AI-TAC in predicting chromatin accessibility of sequences across 81 immune cell types from 6 different lineages[10]. We started with an exploratory analysis to determine the optimal number of units to train ExplaiNN. Saturation in model performance by means of average PCC between predicted and actual ATAC signals, as well as in the number of well-predicted sequences, occurred at ∼250 units (**Fig. 4A**; **Methods**), however, we decided to use 300 units, which is the same number of convolutional filters used in the first layer of AI-TAC. The performance of ExplaiNN by means of average PCC was comparable to that of AI-TAC (1-2% difference; **Fig. 4A**), and the PCCs of individual sequences correlated well between the two models (**Fig. 4B**), however, AI-TAC correctly predicted more sequences (**Fig. 4A**). Next, we visualized both the filters and weights of each unit of the ExplaiNN model, identifying the same lineage-specific TF motifs reported in AI-TAC without having to undergo the computationally intensive process of filter nullification (**Fig. 4C**): NFE2, NFI, and GATA (in stem cells); POU, EBF, and PAX (in B cells); TCF3, TCF7, Ets, and AP1 (in T cells); NR1 (nuclear receptor type 1), TBX, and REL (in innate lymphoid cells); and SPI, Krüppel zinc fingers, and CEBP (in myeloid cells). Moreover, we visualized the importances of each unit to understand their influence on the model’s predictions. For instance, CEBP- and PAX-like units were important for predicting accessibility in most cell types of the myeloid and B lineages, respectively (**Figs. 4D** and **E**). Taken together, using ExplaiNN, we replicated the results of AI-TAC without the need to apply complex and time-consuming interpretation techniques by simply visualizing the weights and importances of each unit.

**Fig. 4:**
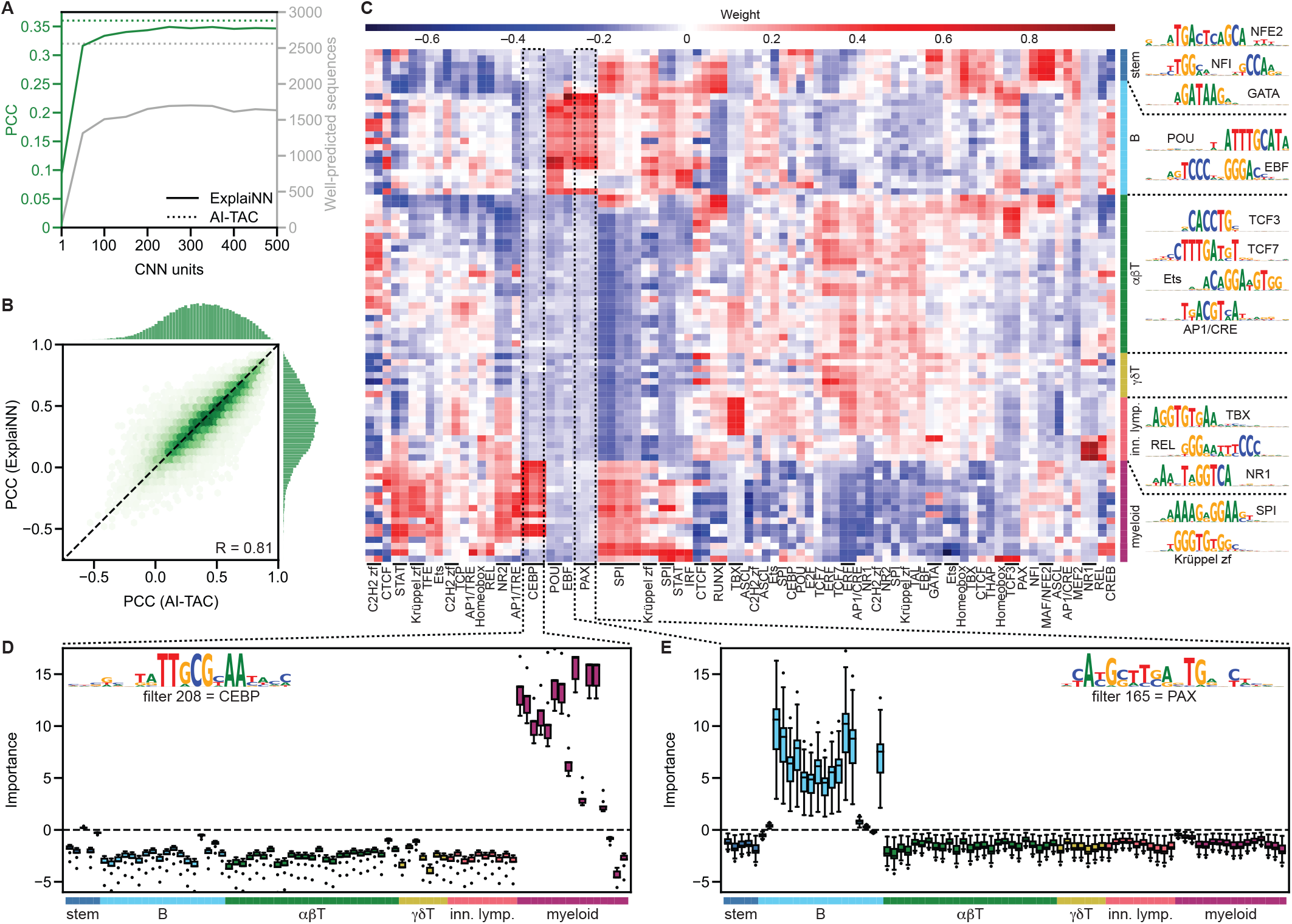
Application of ExplaiNN in predicting chromatin accessibility in the mouse immune system. (**A**) Performances (average PCC; y-axis; green) and number of well-predicted sequences (secondary y-axis; gray) for ExplaiNN models (solid lines) with increasing numbers of units (x-axis) and AI-TAC (dashed lines) trained on OCRs in 81 immune cell types from 6 different lineages[10]. (**B**) Pairwise comparison of the individual performances (PCC) of the OCRs from the previous dataset between the ExplaiNN model trained using 300 units (y-axis) and AI-TAC (x-axis). The Pearson correlation coefficient (R) of the individual OCR performances between the two methods is shown at the lower right corner. (**C**) Heatmap of the final linear layer weights of the ExplaiNN model trained using 300 units, with columns representing units with assigned biological annotations based on their Tomtom[27] similarity to known TF profiles from the JASPAR database[19] and rows representing the 81 immune cell types coloured by lineage: stem cells (navy blue), B cells (turquoise), alpha/beta (forest green) and gamma/delta T cells (olive green), innate lymphoid cells (pink), and myeloid cells (purple). The logos derived from visualizing the filters of selected lineage-specific units are shown at the right. (**D** and **E**) Visualizations of importance scores coloured by lineage for two units from the previous model annotated as CEBP and PAX, revealing their importance to the prediction of chromatin accessibilities in myeloid and B cell lineages, respectively. The logos of the filters of these units are shown at the top. OCR, open chromatin region; PCC, Pearson correlation coefficient; TF, transcription factor.

### ExplaiNN is suitable for the analysis of single-cell chromatin accessibility data

Single-cell (sc) sequencing methods enable profiling of a wide range of genomic information in individual cells (reviewed in [45]), including chromatin accessibility. To explore the utility of ExplaiNN for deciphering *cis*-regulatory properties from granular sc data, we reanalyzed a recent scATAC-seq dataset cataloging 228,873 OCRs across 15,298 human pancreatic islet cells that were grouped into 12 clusters based on their accessibility profiles[46]. We trained a ExplaiNN model with 400 units (*i*.*e*. the number of units at which model performance plateaued) on the sc data to predict the activity of each OCR across the 12 clusters, and visualized both the filters and weights of each unit. We observed that the weights of some units exhibited cell type-specific patterns that had also been found using chromVAR[47] in the original study (**Fig. 5A**). For example, PDX-like units had high positive weights for beta and delta cells, MAF-like units for alpha and beta cells, HNF1-like units for alpha and gamma but not for ductal cells, and FOX-like units for alpha, beta and gamma cells. However, there were also differences: some units did not exhibit high positive weights in expected cell types (*e*.*g*. the highlighted Ets-like unit in endothelial cells), while others exhibited cell type-specific patterns not reported in the original study (**Fig. 5A**). Next, we visualized the importances of each unit for their contributions to the physiological stratification of the cells, finding that RFX-like units were important for predicting OCRs in hormone-high (*i*.*e*. alpha, beta, and delta type 1 cells) but not in hormone-low cells (*i*.*e*. alpha, beta, and delta type 2 cells) (**Fig. 5B**). It was the opposite for AP1-like units: they were important for hormone-low but not hormone-high cells (**Fig. 5C**). Both results were in agreement with the original study. Taken together, ExplaiNN was able to reproduce and expand on the results from the original study while demonstrating its utility for the analysis and interpretation of scATAC-seq data.

**Fig. 5:**
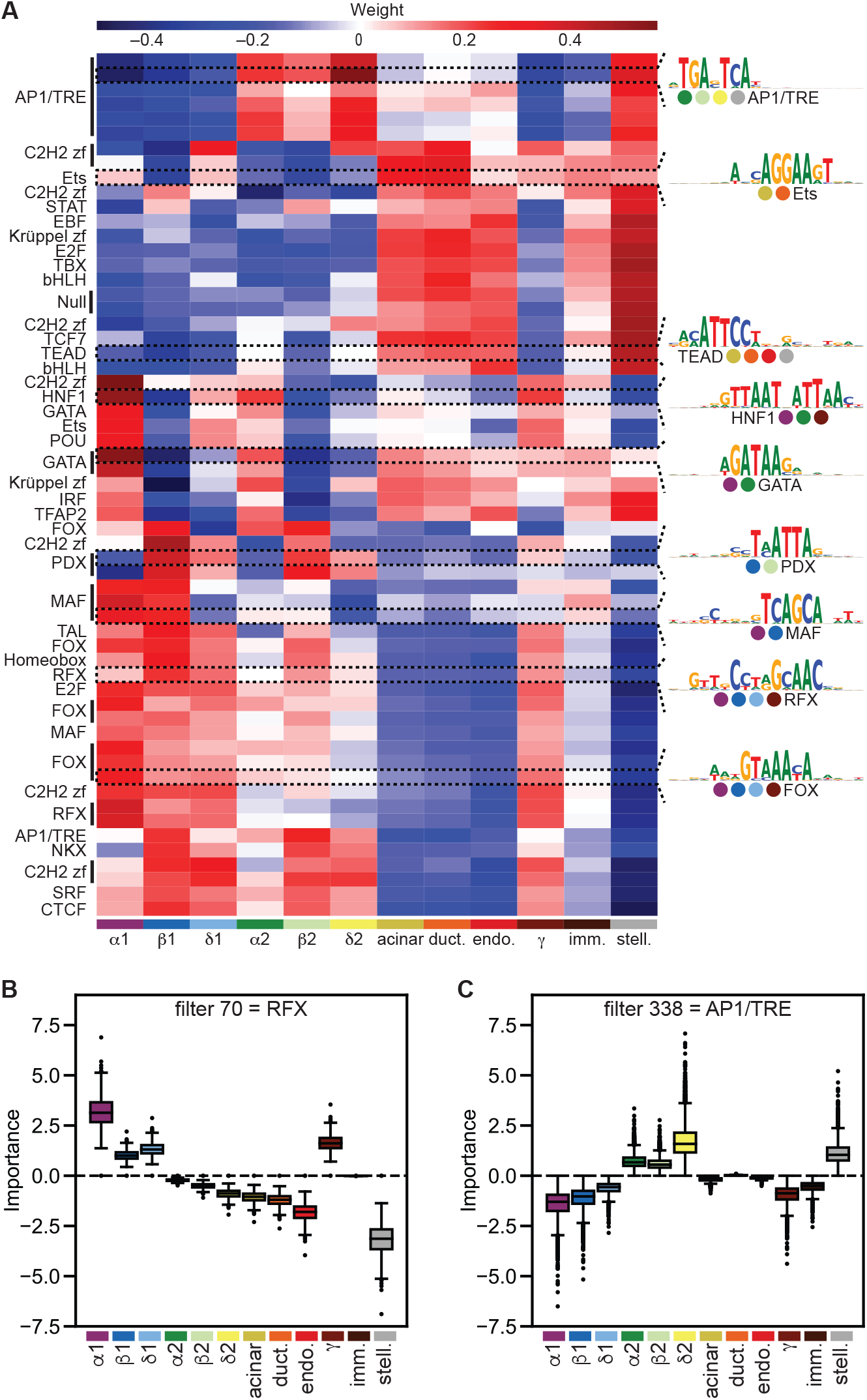
ExplaiNN analysis and interpretation of scATAC-seq data of human pancreatic islets. (**A**) Heatmap of the final linear layer weights of an ExplaiNN model with 400 units trained on OCRs from human pancreatic islet sc data[46], with rows representing units with assigned biological annotations based on their Tomtom[27] similarity to known TF profiles from the JASPAR database[19] and columns representing the 12 clusters of cells based on their accessibility profiles and coloured by cell type: alpha type 1 cells (purple), beta type 1 cells (navy blue), delta type 1 cells (turquoise), alpha type 2 cells (forest green), beta type 2 cells (light green), delta type 2 cells (yellow), acinar cells (olive green), ductal cell (orange), endothelial cells (red), gamma cells (brown), immune cells (dark brown), stellate cells (gray). The logos derived from visualizing the filters of selected units are shown at the right. (**B** and **C**) Visualizations of importance scores coloured by cell cluster for two units from the previous model that were annotated as RFX and AP1 (TRE site), revealing their respective importance to the prediction of chromatin accessibilities in hormone-high and hormone-low cells. OCR, open chromatin region; scATAC-seq, single-cell ATAC-seq; TF, transcription factor.

### ExplaiNN as a plug-and-play platform for TF motifs and deep learning models

Given that ExplaiNN models can be conceptualized as a PWM scanning layer feeding into fully connected layers, we reasoned that initializing the weights of each unit filter with a JASPAR profile would facilitate the interpretation process because the biological annotations of the units would be known beforehand. To confirm this, we trained ExplaiNN models with increasing numbers of units (from 300 to 1,492) on the AI-TAC dataset in which the filter weights of each unit had been initialized with JASPAR profiles (**Fig. 6A**; **Methods**). During training, the filter weights were frozen to prevent them from being refined (*i*.*e*. the models were only allowed to learn the weights of the fully connected and final linear layers). These JASPAR-initialized, frozen models, even the largest one with 1,492 units, performed much worse than both AI-TAC and an ExplaiNN model with 300 units trained from scratch (**Fig. 6B**). Still, the importances of some units were informative. For example, a unit whose filter weights had been initialized with the profile of Lef1 (MA0768.2) was important for predicting accessibility in T cells (**Fig. S4**), in agreement with the role of this TF in establishing T cell identity[48]. We attributed the overall poor performance of these frozen models to the fact that many JASPAR profiles used to initialize the filters might be from TFs irrelevant for immune cells and, therefore, a refinement process would be required to allow them to better resemble the motifs of relevant TFs. Indeed, unfreezing the filter weights during training (and thus allowing their refinement) improved the performance of the model, approaching that of AI-TAC (**Fig. 6B**). Moreover, as a consequence of this refinement, >35% of the filters underwent substantial changes; their visualization as PWMs revealed that they had become different from the original JASPAR profiles used for their initialization (Tomtom[27] *q*-value >0.05). For instance, a unit whose filter weights had been initialized with the profile of TFAP2C (MA0815.1), and whose importance across the different immune lineages when freezing the filter weights during training was null, became important for predicting accessibility in B cells; its filter was refined to such an extent that when visualized as a PWM it resembled the motif of EBF1, an important TF for maintaining B cell identity[49] (**Fig. 6C**).

**Fig. 6:**
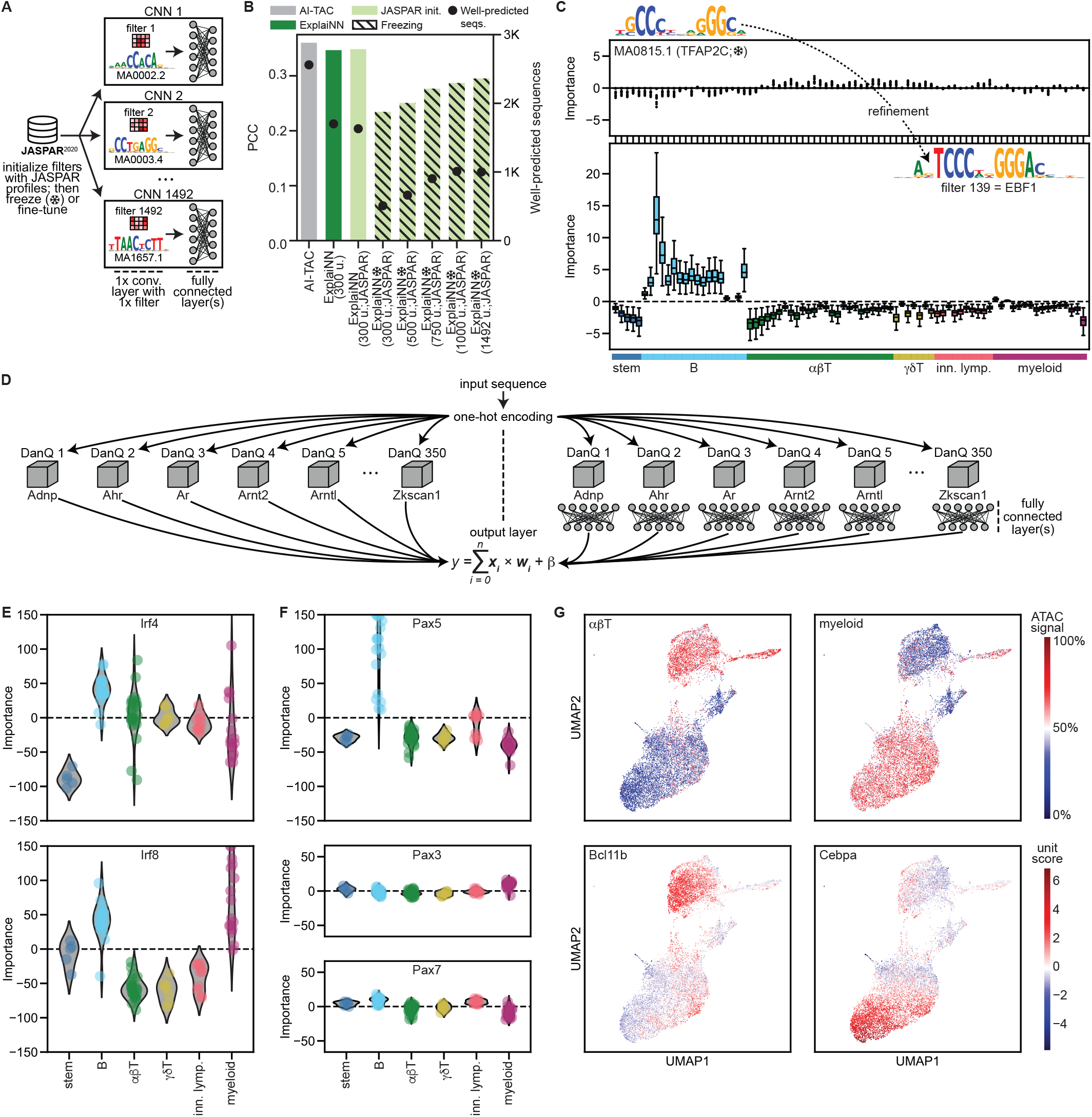
Initializing ExplaiNN with JASPAR profiles and DanQ models. (**A**) Schematic representation of the proposed ExplaiNN model in which the convolutional filters of each unit are initialized with JASPAR[19] profiles. (**B**) Performance (average PCC; y-axis) and number of well-predicted sequences (secondary y-axis; black dots) for different models trained on OCRs from the AI-TAC dataset[10]: one ExplaiNN model with 300 units that was trained normally (green), as well as different ExplaiNN models with increasing numbers of units (300, 500, 750, 1000, 1492) that were initialized with JASPAR profiles (light green), with (pinstripes) and without (no pinstripes) freezing the weights of the filters during training. The performance of AI-TAC is provided as baseline (gray). (**C**) Refinement example for a filter whose weights had been initialized with the JASPAR profile of TFAP2C (MA0815.1). Freezing the filter weights during training was detrimental (top): the importance scores of that filter’s unit across all immune lineages was null. However, if filter weight refinement was allowed during training (*i*.*e*. no freezing; bottom), that filter was modified until it resembled the motif of EBF1, an important TF for maintaining B cell identity[49], and that same unit became important for predicting accessibility in B cells. (**D**) Schematic representation of the proposed transfer learning strategy in which ExplaiNN units are replaced with pretrained DanQ[8] models that can be followed (right) or not (left) by fully connected layers. (**E** and **F**) We used the transfer learning strategy on the left to train one ExplaiNN model with 350 units on the AI-TAC dataset in which the units had been replaced with 350 different pretrained DanQ models, each predicting the binding of a single TF to the mouse genome. During the training process of the ExplaiNN model, the DanQ models were frozen (*i*.*e*. their weights were not modified). Unit importance scores of DanQ models belonging to different members of the IRF and PAX TF families are shown. (**G**) We repeated the same process but using the transfer learning strategy on the right: each unit was replaced with a pretrained DanQ model but adding two fully connected layers after each model. Then, we applied UMAP[55] to cluster the OCRs based on their unit outputs. (top) UMAP clusters display cell-type specificity (*e*.*g*. alpha/beta T and myeloid cells). (bottom) The outputs of the Bcl11b and Cebpa DanQ model units strongly agree with their biologically relevant clusters. OCR, open chromatin region; PCC, Pearson correlation coefficient; TF, transcription factor; UMAP, uniform manifold approximation and projection.

Another limitation that arises during model interpretation is the inability to distinguish between TFs from the same structural family (*i*.*e*. TFs sharing the same class of DNA-binding domain), as they often have highly similar DNA-binding specificities[50]. In order to generate individual units with the resolution to highlight the importance of different paralogs, we implemented a transfer learning strategy (**Fig. 6D**; **Methods**): We pretrained 350 single-task DanQ models, each predicting the binding of a single TF to the mouse genome. The AUPRCs of the pretrained DanQ models ranged from 0.51 (E4f1) to 0.96 (Snai2), with a median of 0.78 across all models (**Table S2**). Then, we initialized an ExplaiNN model with 350 units in which we replaced the layers of each unit with one of the pretrained DanQ models. Similar to NAMs, we initialized a second ExplaiNN model in which we added two fully connected layers after each DanQ model. Next, we fine-tuned both ExplaiNN models on the AI-TAC dataset, freezing the pretrained DanQ models (*i*.*e*. their weights were not modified). The performance of the fine-tuned models nearly reached that of AI-TAC (PCC = 0.335 and 0.343 vs. 0.360 for AI-TAC). Next, we visualized the importances of each DanQ model unit in the ExplaiNN model without fully connected layers, allowing us to disambiguate the contribution of individual TF family members (**Methods**). For instance, the Irf4 and Irf8 units were important for predicting accessibility in different immune lineages (**Fig. 6E**), in agreement with the distinct roles of these TFs: Irf4 regulates B, T, myeloid, and dendritic cell differentiation[51], while Irf8 regulates B and myeloid cell lineages[52]. Similarly, the Pax5 unit was important for predicting accessibility in B cells, consistent with the role of this TF in establishing B lineage identity and function[53]; in contrast, the importances of the Pax3 and Pax7 units across immune lineages were negligible, in agreement with their role in regulating myogenesis[54] (**Fig. 6F**). Finally, we applied UMAP[55] to cluster the sequences based on their unit outputs in the second ExplaiNN model, resulting in three main clusters that were associated with ATAC signals in alpha/beta T, myeloid, and B cells (**Figs. 6G** and **S5**). The outputs of some units were in strong agreement with a biologically relevant cluster. For example, the Bcl11b unit outputs were confined within the boundaries of the alpha/beta T cell cluster, consistent with its role in the differentiation and survival of these lymphocytes[56], the Cebpa unit outputs within the myeloid cluster, in agreement with its role in myeloid differentiation[57], and the Ebf1 unit outputs within the B cluster (**Figs. 6G** and **S5**). Taken together, replacing the units of ExplaiNN with pretrained high-resolution TF binding models achieved performance levels comparable to state-of-the-art deep learning models while providing the additional value of gaining insights into the roles of individual TFs. This flexibility of incorporating different components into ExplaiNN offers the potential for increased performance while retaining interpretability.

## Discussion

ExplaiNN is an explainable deep learning approach designed for straightforward discovery of genomic features contributing to model performance for a broad range of predictive tasks related to DNA sequence data. Inspired by the recently introduced NAMs method, the architecture of ExplaiNN is based on multiple, simple, independent CNN units that recognize sequence patterns. The outputs of these units are subsequently combined in a similar manner to classical regression analysis techniques. We benchmarked ExplaiNN on diverse tasks, including sequence-based prediction of TF binding, chromatin accessibility, both from bulk and single-cell data, enhancer activity, and *de novo* motif discovery. Through this array of diverse applications, ExplaiNN performed comparably to leading methods specialized for the independent tasks.

While deep learning models have been successful at revealing novel biological insights from high-throughput sequencing data, outperforming traditional PWM-based approaches such as motif enrichment analysis, their adoption in the genomics field remains largely in the hands of deep learning experts. This is in part a consequence of their perceived opaque nature and lack of consensus on best practices for their interpretation, making it difficult for the non-specialist to apply these methods routinely. There is a growing need for deep learning models that are more interpretable and easy to use[15,16]. ExplaiNN is one such approach. Architecturally, ExplaiNN models are constrained to only capturing homotypic cooperativity, excluding heterotypic interactions between pairs of motifs. It may therefore be surprising to some that in practice ExplaiNN performs so well across a wide range of predictive tasks. One reason behind this could be that excluding higher-order interactions acts as a regularizer leading to more accurate predictions than multi-layer CNN models on simple tasks such as TF binding prediction. Another reason could be that the presence of higher-order interactions in the genomics explored datasets could be marginal, in agreement with recent studies suggesting that *cis*-regulatory logic may be less complex than some believe[58,59]. Alternatively, these kinds of complex interactions may be highly specific to individual enhancers, and thus not detectable by models generalized for analysis across the genome. As exemplified by the DeepSTARR analysis in this report, comparisons between ExplaiNN and non-linear models can enable researchers to assess the benefit from allowing non-linear interactions. In such cases where there is not substantial benefit, the use of the simpler, readily interpretable method may be preferred.

The emergence of new methods for regulatory sequence analysis is a recurrent process. An important early approach to identifying recurring subsequences associated with TFBSs used the expectation-maximization algorithm to learn a PWM from a set of unaligned regulatory sequences[60]. These individually learned PWMs could then be used in a linear model to make predictions regarding various properties of a given sequence[61]. ExplaiNN could be perceived as a direct extension of this approach, wherein its convolutional filters learn a large amount of PWMs simultaneously, however, unless the fully connected layers of each unit were replaced with a global maximum pooling layer, their activation values would not be equivalent to those of independent PWM scans. The fully connected layers might transform the activation values into more expressive and informative scores leading to better performance. As we have shown, this expressiveness is likely achieved by learning relevant motif positional information.

As shown throughout this work, ExplaiNN has the potential to be applied to multiple problems. This flexibility may motivate users to become proficient with the system as they will be able to work across or within topics using the same tool. As presented here for *de novo* motif discovery, the flexibility to apply ExplaiNN to both ChIP-seq data and multiple special classes of *in vitro* TF binding data has the promise to reduce the number of methods that must be mastered.

There remains an ample variety of potential directions for further development of ExplaiNN, split between enhanced technical capabilities and user empowerment. It should be possible to integrate a more complex model architecture within the system to account for the “missing complexity” (*e*.*g*. non-linearity). As implemented, ExplaiNN cannot address analyses requiring large receptive fields (*e*.*g*. Akita[6], Xpresso[12], Basenji[62] or Enformer[13]), but it should be possible to implement such capacity within the system. Incorporating more advanced visualization techniques and data processing tools would ease the adoption of ExplaiNN by scientists working on applied problems who currently rely on heritage methods for regulatory sequence analysis.

The software for using ExplaiNN is provided open-source in a well-documented repository, and it is our hope that it will lead to or inspire widespread use of interpretable deep learning methods.

## Methods

### Model architectures

Unless otherwise specified, each ExplaiNN unit consisted of:

- 1st convolutional layer with 1 filter (19×4), batch normalization, exponential activation and max pooling (kernel size=7, stride=7);
- 1st fully connected layer with 100 nodes, batch normalization, ReLU activation and 30% dropout; and
- 2nd fully connected layer with 1 node, batch normalization and ReLU activation.

In the 1st convolutional layer, exponential activation was used (as opposed to ReLU), as it has been shown to significantly improve the recovery of biologically meaningful motifs from the filters[26]. The final layer of ExplaiNN is linear: one fully connected output layer with *n* outputs (*e*.*g*. 50 in the initial TF binding prediction task or 81 when predicting chromatin accessibility states across immune cells).

For the CNN_1_ and CNN_1_Exp architectures, we used the following specifications:

- 1st convolutional layer with 100 filters (19×4), batch normalization, ReLU (for CNN_1_) or exponential (for CNN_1_Exp) activation and max pooling (kernel size=7, stride=7);
- 1st fully connected layer with 1000 nodes, batch normalization, ReLU activation and 30% dropout;
- 2nd fully connected layer with 1000 nodes, batch normalization, ReLU activation and 30% dropout; and
- Fully connected output layer with 50 outputs.

For the DeepCNN (adapted from Basset[9]):

- 1st convolutional layer with 100 filters (19×4), batch normalization, ReLU activation and max pooling (kernel size=3, stride=3);
- 2nd convolutional layer with 200 filters (7×1), batch normalization, ReLU activation, and max pooling (kernel size=3, stride=3);
- 3rd convolutional layer with 200 filters (4×1), batch normalization, ReLU activation, and max pooling (kernel size=3, stride=3);
- 1st fully connected layer with 1000 nodes, batch normalization, ReLU activation and 30% dropout;
- 2nd fully connected layer with 1000 nodes, batch normalization, ReLU activation and 30% dropout; and
- Fully connected output layer with 50 outputs.

For DanQ[8]:

- 1st convolutional layer with 320 filters (26×4), ReLU activation, 20% dropout, and max pooling (kernel size=13, stride=13);
- 2 bi-directional LSTM layers with hidden state size 320 and 50% dropout;
- 1st fully connected layer with 925 nodes and ReLU activation; and
- Fully connected output layer with 1 or 50 outputs (depending on the task).

For DeepSTARR[30]:

- 1st convolutional layer with 256 filters (7×4), padding of size 3, batch normalization, ReLU activation function and max pooling (kernel size=2, stride=2);
- 2nd convolutional layer with 60 filters (3×1), padding of size 1, batch normalization, ReLU activation function and max pooling (kernel size=2, stride=2);
- 3rd convolutional layer with 60 filters (5×1), padding of size 2, batch normalization, ReLU activation function and max pooling (kernel size=2, stride=2);
- 4th convolutional layer with 120 filters (3×1), padding of size 1, batch normalization, ReLU activation function and max pooling (kernel size=2, stride=2);
- 1st fully connected layer with 256 nodes, batch normalization, ReLU activation function and 40% dropout;
- 2nd fully connected layer with 256 nodes, batch normalization, ReLU activation function and 40% dropout; and
- Fully connected output layer with 2 outputs.

The AI-TAC architecture was provided by Maslova and colleagues[10]. All architectures were implemented using the PyTorch framework[63] (version 1.9.0).

### Interpretability with ExplaiNN

First, we converted the filter of each unit into a PWM by following the specifications from the AI-TAC manuscript: For each filter, we built a position frequency matrix (PFM) by aligning all 19-mers (*i*.*e*. 19 bp-long DNA sequences) activating that filter’s unit by ≥50% of its maximum activation value in correctly predicted sequences. The resulting PFM was then transformed into a PWM by setting the background uniform nucleotide frequency to 0.25. Next, the PWM derived from each filter was mapped to one or more profiles from the vertebrate collection of the JASPAR 2020 database[19] using Tomtom[27] (version 5.3.0; q-value ≤0.05), which, in turn, were used to annotate that filter’s unit. For instance, a unit whose filter’s PWM were similar to the profiles of members from the SOX TF family would be annotated as “SOX-like”.

For global interpretability, we visualized the weights of each unit in the final linear layer of the model (*e*.*g*. using a heatmap). Usually, these weights were associated with the importance of the unit for each task. In some cases, however, they were highly positive or negative for a task in which the unit was not activated by the input sequences. To overcome this limitation and obtain more fine-grained unit importances, for each unit and for each task, we computed the product of the unit’s activation for each sequence with the final layer weight for that task, and visualized them (*e*.*g*. using a boxplot). For visualization, for each unit, we only included the products from correctly predicted sequences activating that unit’s filter by ≥50% of its maximum activation value. In addition, for each unit and for each task, the median of these products was used to assess the importance of that unit for that task.

### Interpretability with DeepLIFT and TF-MoDISco

For each correctly predicted sequence in the test set, we generated DeepLIFT[23] importance scores with 10 reference sequences using the Captum library[64] (version 0.4.0). We used TF-MoDISco[24] (version 0.5.14.0) with default settings to obtain motifs from DeepLIFT importance scores.

### Training, validation and test datasets

To obtain human *in vivo* TF binding data, we repurposed a previously described data matrix aggregating the binding of 163 TFs to 1,817,918 200-bp-long DNase I hypersensitive sites (DHSs) in 52 cell and tissue types[65]. For each TF-DHS pair, a “1” was used to indicate that the DHS was accessible and bound by the TF (*i*.*e*. the DHS and at least one ChIP-seq peak summit of that TF from ReMap 2018[44] overlapped), a “0” that the DHS was accessible but not bound by the TF, and a null sign (“∅”) that it was not accessible to the TF for binding (*i*.*e*. unresolved). For the CNN_1_, CNN_1_Exp, DeepCNN, DanQ, and ExplaiNN models predicting the binding of 50 TFs, we extracted a slice of the matrix including the row vectors of the 50 TFs, removing any column vectors with unresolved elements. The resulting resolved regions for all 50 TFs were randomly split into training (80%), validation (10%) and test (10%) sets using the “train_test_split” function from scikit-learn[66] (version 0.24.2; datasets were always randomly split in this way). For *de novo* motif discovery, since the number of bound versus unbound regions of all TFs were imbalanced, we subsampled the set of unbound regions to a 50:50 ratio for each TF while accounting for their %GC content distributions. The resulting datasets were randomly split into training (80%), validation (10%) and test (10%) sets while maintaining an equal proportion of bound and unbound regions.

Mouse *in vivo* TF binding data was obtained as follows: First, we downloaded non-redundant mouse ChIP-seq peaks for 350 DNA-binding TFs[67] from ReMap. We resized each ChIP-seq peak to 201 bp by extending its summit 100 bp in both directions. Furthermore, we retrieved an atlas of 1,802,604 DHS regions in the mouse genome[68], which we also resized to 201 bp around the center of each DHS. In both cases, we applied BEDTools slop[69] (version 2.30.0). We then created a set of negative sequences for each TF by subsampling non-overlapping regions from the DHS atlas while matching the %GC content distribution of its ChIP-seq peaks. The resulting sequences for each TF were randomly split into training (80%), validation (10%) and test (10%) sets while keeping the same ratio between positive and negative sequences.

PBM data was downloaded from UniPROBE[40] (Gata3; UP00032; clone ID pTH1024). Probe signal intensities were quantile normalized (QN) using the “quantile_transform” function from scikit-learn. Probes were 60 bp long, including both the de Bruijn and linker sequences. The arrays AMADID #015681 and #016060 were used for training and validation, respectively.

HT-SELEX[39] and SMiLE-seq[41] data were retrieved from the Sequence Repository Archive (run ids: ERR1003435, ERR1003437, ERR1003439, ERR1003441, and SRR3405148) using parallel-fastq-dump (version 0.6.7). For HT-SELEX data, we treated each cycle as an independent class as in Asif and Orenstein[70], thereby removing the need for negative sequences. Reads were randomly split into training (80%) and validation (20%) sets while preserving the proportions between reads from each cycle. For SMiLE-seq data, reads were left- and right-clipped 7 and 64 bp, respectively, for a final length of 30 bp corresponding to the randomized DNA. A set of negative sequences were obtained by dinucleotide shuffling using BiasAway[71] (version 3.3.0). Sequences were randomly split into training (80%) and validation (20%) while maintaining an equal proportion between positives and negatives.

Binarised human pancreatic islet scATAC-seq data was obtained from [46]. Data was denoised by training a PeakVI[72] model with 21 latent dimensions using scvi-tools[73] (version 0.14.6). The denoised profiles were used to compute the mean accessibility of each peak in the 12 clusters identified in the original study. Peaks were resized to 600 bp around the center of each peak using BEDTools slop. Peaks from chromosome 1 were used for validation, peaks from chromosomes 10, 11, and 12 for testing, and peaks from the remaining chromosomes for training.

The AI-TAC and DeepSTARR datasets were obtained from the original publications, and we used the same data splits.

### Model training

Unless otherwise specified, we trained all models using the Adam optimizer[74] and binary cross entropy as loss function, except for models trained on PBM data, wherein we used the mean squared error. We applied one-hot encoding to convert nucleotides into 4-element vectors (*i*.*e*. A, C, G, and T), set the learning rate to 0.003 and batch size to 100, and used an early stopping criteria to prevent overfitting when the model performance on the validation set did not improve. The training and validation sets included the reverse-complement of each sequence. Sequences with Ns were discarded.

### *De novo* PWMs

For each of the 163 TFs from the data matrix, we trained one ExplaiNN model with 100 units using its training and validation sets, and then visualized the filters and importance scores of each unit, resulting in 100 PWMs per TF. As baseline, we applied STREME[34] (version 5.3.0) on the set of training sequences of each TF to also generate 100 PWMs: We set the fraction of sequences held-out for *p*-value computation to 0 (option --hofract), the maximum length of the PWMs to 19 bp (*i*.*e*. the filter size in ExplaiNN; option --maxw), and the number of output PWMs to 100 (option --nmotifs). Next, for each TF and for each of its PWMs, we evaluated the PWM performance by keeping the score of the best hit from scanning that PWM along both strands of each sequence in the test set and computing the AUPRC.

Like above, for each GATA3 *in vitro* assay (*i*.*e*. HT-SELEX, PBM and SMiLE-seq), we trained one ExplaiNN model with 100 units using the training and validation sets, and visualized the filter of each unit, resulting in 100 PWMs for each assay. However, the task of the ExplaiNN model trained on HT-SELEX data was multiclass classification: the model tried to predict the cycle of origin of each input read, with the expectation that the last cycles would be enriched for bound sequences of the TF. In addition, for PBM data, the task of the ExplaiNN model was to infer QN intensity signals of each probe.

For each TF and for each assay, the best PWM was selected based on its performance on the corresponding validation set.

### Plug-and-play

First, we resized all 746 profiles from the JASPAR 2020 vertebrate collection to 19 bp. Then, we applied a farthest point sampling procedure to remove profile redundancy based on Tomtom similarities starting from the most dissimilar profile. This resulted in four different sets of non-redundant profiles of sizes 150, 250, 375 and 500. Next, we computed the reverse complement of each profile, doubling the number of profiles in each non-redundant set to 300, 500, 750 and 1000, and raising the number of profiles in the JASPAR 2020 vertebrate collection to 1492. We trained ExplaiNN models with increasing numbers of units (300, 500, 750, 1000, and 1492) on the AI-TAC dataset in which the filter weights of each unit were initialized with those JASPAR profiles. To initialize filter weights with profiles from JASPAR, we followed the specifications from DanQ: We reformatted JASPAR profiles as PWMs using Biopython[75] (version 1.79) and then converted them to filter weights by subtracting 0.25 from the probability of each nucleotide at each PWM position. During training, nullification of gradients could be applied to freeze the filter weights.

For each of the 350 TFs from the mouse *in vivo* TF binding data, we trained a DanQ model using the training and validation sets of that TF, and assessed its performance by computing the AUPRC on the test set. We then scored the sequences from the AI-TAC dataset keeping the outputs from each model (pre- or post-sigmoid). Next, we combined the outputs of each sequence in one fully connected layer with 81 output nodes. Alternatively, each DanQ output could be embedded in its own fully connected neural network with one layer and 200 nodes, resulting in 350 processed outputs ultimately combined in one fully connected layer with 81 outputs.

UMAP clusters were obtained and plotted using the UMAP Python library[55] (version 0.5.2).

## Code availability

ExplaiNN is open-source software distributed under the MIT license. It is available on GitHub (https://github.com/wassermanlab/ExplaiNN) accompanied by the IPython notebooks that we used for the different experiments, as well as a collection of Python scripts to allow users to deploy their own ExplaiNN models.

## Data availability

The TF binding matrices, and the code for generating them, are available on GitHub as 2D numpy arrays[76] (https://github.com/wassermanlab/TF-Binding-Matrix). The filter weights from JASPAR profiles, and the code for generating them, are also available on GitHub as pickles (https://github.com/wassermanlab/PWM-to-filter-weights). The remaining datasets used in this work were obtained either from the corresponding manuscripts or from specialized repositories, as indicated in the **Methods** section.

## Author contributions

MS and GN conceived the idea; OF, GN and MS designed and performed the analyses; GN, MS and OF wrote the code; GN developed the Python library; OF and GN created the figures with input from MS; WW and SM provided advice on all aspects of the work; all authors participated in the writing and editing of the manuscript.

## Acknowledgements

We thank the members of the Mostafavi and Wasserman labs for providing useful feedback. We thank Dora Pak and Jonathan Chang for administrative and IT support, respectively. We thank UBC’s Advanced Research Computing (ARC) (https://arc.ubc.ca/) for enabling HPC.

## Funding

GN was supported by an International Doctoral Fellowship from the University of British Columbia. OF, MS and WWW were supported by grants from the Canadian Institutes of Health Research (PJT-162120), Natural Sciences and Engineering Research Council of Canada (NSERC) Discovery Grant (RGPIN-2017-06824), and the BC Children’s Hospital Foundation and Research Institute. SM acknowledges support from the Canadian Institute for Advanced Research (CIFAR). Equipment enabled by NSERC Research Tools and Instruments Grant (RTI-2020-00778) to SM and WWW.

## Figures

**Fig. S1:** From top to bottom, visualization of importance scores for three units annotated as CTCF, one unit annotated as CEBP, and one unit annotated as Ets from an ExplaiNN model trained using 100 units on predicting the binding of 50 TFs in OCRs. OCR, open chromatin region.

**Fig. S2:** Cooperativity (residual fold change; y-axis) plotted as a function of distance (x-axis) between the motifs of the housekeeping TFs Dref (top row, yellow), Ohler1 (middle row; blue), and Ohler6 (bottom row; green) for an ExplaiNN model in which the fully connected layers of each unit had been replaced with a global max pooling layer. The 5-mer GGGCT is provided as a negative control (light gray). TF, transcription factor.

**Fig. S3:** Logos derived using ExplaiNN or STREME[34] for the nuclear receptors ESR1, ESRRA, HNF4A, HNF4G, NR2C2, NR2F1, NR2F6, NR3C1, and RXRA on *in vivo* datasets describing the binding of these TFs to OCRs. For comparison, the JASPAR[19] logos for these TF profiles are shown: MA0112.3 (ESR1), MA0592.3 (ESRRA), MA0114.4 and MA1494.1 (HNF4A), MA0484.2 (HNF4G), MA0504.1 and MA1536.1 (NR2C2), MA0017.2 and MA1537.1 (NR2F1), MA0677.1 (NR2F6), MA0113.3 (NR3C1), and MA0512.2 (RXRA).

**Fig. S4:** Visualization of importance scores coloured by lineage of a unit from an ExplaiNN model initialized with 300 JASPAR profiles and trained on the AI-TAC dataset[10] with freezing. The unit, which corresponded to the JASPAR profile of Lef1 (MA0768.2), was important for predicting accessibility in T cells, in agreement with the role of this TF in establishing T cell identity[48]. TF, transcription factor.

**Fig. S5:** ExplaiNN model with 350 units trained on the AI-TAC dataset[10] in which the units had been replaced with 350 different pretrained DanQ[8] models, each predicting the binding of a single TF to the mouse genome. During the training process of the ExplaiNN model, the DanQ models were frozen (*i*.*e*. their weights were not modified). TF, transcription factor.

## Tables

**Table S1:** List of TF binding modes used in this study.

**Table S2:** Individual performances of 350 DanQ models trained on mouse *in vivo* binding data in predicting DNA binding of their respective TFs.

## Notes

https://github.com/wassermanlab/ExplaiNN

### Competing Interest Statement

The authors have declared no competing interest.

### Summary of Updates

We have added a new figure (new Fig. 2) to show that ExplaiNN can capture homotypic cooperative interactions, as well as a new panel to Fig. 3 and a new supplementary figure (new Fig. S3) to demonstrate its capacity to learn full motif representations.

